# Transcriptomic responses and survival mechanisms of staphylococci to the antimicrobial skin lipid sphingosine

**DOI:** 10.1101/2020.09.15.297481

**Authors:** Yiyun Chen, Josephine C. Moran, Stuart Campbell-Lee, Malcolm J. Horsburgh

## Abstract

Sphingosines are antimicrobial lipids that form part of the innate barrier to skin colonisation by microbes. Sphingosine deficiencies can result in increased epithelial infections by bacteria including *Staphylococcus aureus*. Recent studies have focused on the potential use of sphingosines as novel therapeutic agents, but there have been no investigations into sphingosine resistance or its potential mechanisms. We used RNA-Seq to identify the common D-sphingosine transcriptomic response of the transient skin coloniser *S. aureus* and the dominant skin coloniser *S. epidermidis*. A common D-sphingosine stimulon was identified that included downregulation of the SaeSR two-component system (TCS) regulon and upregulation of both the VraSR TCS and CtsR stress regulons. We show that the PstSCAB phosphate transporter, and VraSR offer intrinsic resistance to D-sphingosine. Further, we demonstrate increased sphingosine resistance in these staphylococci evolves readily through mutations in genes encoding the FarE-FarR efflux/regulator proteins. The ease of selecting mutants with resistance to sphingosine may impact upon staphylococcal colonisation of skin where the lipid is present and have implications with topical therapeutic applications.

## Introduction

Human skin is a protective barrier with lipids that contribute a major role in establishing the innate defensive properties. Skin lipids of human epidermis have two distinct origins: sebaceous glands, and keratinocytes of the stratum corneum. Composition of these two lipid fractions differ markedly with sebaceous lipids being predominantly squalene, wax esters and triacyclglycerols, whereas mostly the epidermal lipids comprise equimolar amounts of ceramides, free fatty acids and cholesterol (Nicolaides, 1974, Pappas, 2009, Coates et al., 2014). The major antimicrobial lipids of epidermal and sebaceous fractions also differ; both contain antimicrobial fatty acids, but the hydrolysis of ceramides generates antimicrobial sphingosines only in the epidermal lipids fraction.

Sphingosines are unsaturated long chain amino alcohols and their rapid antimicrobial activity contributes to growth limitation of skin pathogens, such as *Staphylococcus aureus* (Bibel et al., 1992, Parsons et al., 2012). *S. epidermidis* is a dominant skin coloniser but the effects of sphingosines upon their colonisation are unknown. Reduced sphingosine concentration was correlated with increased skin colonisation by *S. aureus* in patients with atopic dermatitis (Arikawa et al., 2002), and with *S. aureus* lung infections in cystic fibrosis patients (Tavakoli Tabazavareh et al., 2016). The concentration of sphingosines is significantly reduced in the lungs of cystic fibrosis (CF) patients due to lower ceramidase activity, and in CF mouse models (Pewzner-Jung et al., 2014), which are both more susceptible to pulmonary infections. The increased susceptibility to *S. aureus* due to the imbalance between ceramides and sphingosines is reversible with sphingosine treatment in mouse models (Tavakoli Tabazavareh et al., 2016). Sphingosine was proposed as a novel treatment against staphylococcal infection; as a pre-treatment on implanted medical devices (Seitz et al., 2019, Beck et al., 2020), or as an inhaled treatment for pulmonary *S. aureus* infections in CF patients (Tavakoli Tabazavareh et al., 2016, Verhaegh et al., 2020).

Sphingosine was recently shown to interact with the cell membrane via cardiolipin to kill *S. aureus* through rapid membrane permeabilization (Verhaegh et al., 2020). The authors speculate that cardiolipin-sphingosine interaction forms rigid domains within the membrane, as described for the aminoglycoside 3’, 6-dinonylneamine (Verhaegh et al., 2020, El Khoury et al., 2017). Knowledge of resistance mechanisms within staphylococcal communities would help to inform usage and to safeguard against rapid spread of resistance to this novel antimicrobial.

In this study, we compared the gene expression of both *S. aureus* and *S. epidermidis* in response to micromolar concentrations of D-sphingosine and identified a common stimulon. Discrete responses were revealed, and a common mode of resistance was identified through experimental evolution of both species of staphylococci.

## Materials and Methods

### Bacterial strains and culture

Staphylococci used in this study included the *S. aureus* strains: SH1000, Newman, SF8300, MRSA252, MSSA476 and BL137; and *S. epidermidis:* Tü3298, NCTC 1457, Rp62a, B19 and O16. Construction of a *farE::tet* strain of *S. aureus* SH1000 and its complementation was described in Kenny et al. (2009) as *SAR2632::tet*. Strain SH1000 FarR.D94Y.E151K was constructed with plasmid pMutin4 by allelic replacement as previously described (Horsburgh et al., 2004) using primers *farR_HindIII* and *farR_BamHI* (Table S1) to PCR amplify and clone the mutant allele. The mutations from the experimentally evolved strain were recombined into SH1000 and resolved for loss of the plasmid, with correct construction of SH1000 FarR.D94Y.E151K confirmed using Sanger sequencing. Mutants *vraS::tet* and *vraR::tet* were constructed previously (Kenny et al., 2009).

An operon deletion mutant of *pstSCAB::tet* was generated in *S. aureus* Newman using pMutin4 by cloning partial *pstS* and *pstB* gene fragments together with the tetracycline resistance cassette, as described in Kenny *et al*. (2009). Mariner transposon mutants of *pstS, phoR, hrtB* and *phoU* from the Nebraska Transposon library of *S. aureus* USA300 were transduced to strain Newman, as described previously (Fey et al., 2013, Horsburgh et al., 2001), using gene-specific primers paired with described upstream/buster primers to confirm orientation and verify their correct allelic replacement; gene-specific primers were used for mutants of: *phoU, phoR, hrtB* and *pstS* (Table S1). D-sphingosine (Sigma-Aldrich) stock solution was prepared at a concentration of 5 mg ml^-1^ in ethanol. Antibiotics tetracycline, erythromycin, lincomycin and ampicillin were used at concentrations 5 μg ml^-1^, 5 μg ml^-1^, 25 μg ml^-1^ and 100 μg ml^-1^ respectively.

### Minimum inhibitory concentration (MIC) assay

MIC assay used a broth microdilution method with cultures inoculated at OD_600_ 0.1 and growth assessed after 24 h incubation at 37 °C.

### Sphingosine challenge for RNA analysis

Bacterial challenge growth experiments were conducted as described previously (Moran et al., 2017), with the exception that D-sphingosine was added to the culture for the challenge at a concentration of 5 μM.

### RNA and DNA sequencing and bioinformatic analysis

RNA extraction used enzymatic lysis as described previously (Moran et al., 2017), with key features being that RNA was rRNA depleted using RiboZero and libraries constructed using ScriptSeq (Epicentre). Samples were sequenced by paired-end sequencing on the HiSeq platform (Illumina). RNA from the same samples that was not used for sequencing was converted to cDNA and used for qPCR assays. Bowtie 2 (Langmead and Salzberg, 2012) was used to map reads and Edge R (Robinson et al., 2010, McCarthy et al., 2012) was used to determine differentially expressed genes. Benjamin and Hochberg analysis false discovery rate <0.05 was used as the cut off for differentially expressed genes between control and test conditions. RNA sequencing and analysis was undertaken by the Centre for Genomic Research (CGR, Liverpool).

For the sequencing of experimentally evolved isolates, DNA was extracted, sequenced and analysed as described previously (Coates-Brown et al., 2018), with the exception that all isolates were sequenced individually rather than pooled. Mapping statistics are included as Table S2.

### cDNA generation and qPCR conditions

cDNA was synthesized from high integrity RNA samples using the tetro cDNA synthesis kit (Bioline). The SensiFAST SYBR Hi-ROX kit (Bioline) was used for qPCR reactions. The reaction mix contained 10 μl Sensifast, 0.5 μM of each primer, 100 μg cDNA wth DEPC-treated water in a reaction volume of 20 μl. At least three biological replicates were used. Primers used for qPCR are listed in Table S1.

### Staphyloxanthin extraction assay

Bacteria were grown under challenge growth conditions. At 25 h cells were adjusted to OD_600_=5, and 10 ml harvested by centrifugation then resuspended in 1 ml methanol and incubated for 15 min at 37°C with shaking. The suspension was centrifuged for 15 min at 5k rpm and using 250 μl of supernatant, absorbance was assayed in triplicate for across wavelengths 350-550 nm.

### Sphingosine challenge survival assay

Bacteria were cultured overnight at 37°C in the absence or presence of 5 μM D-sphingosine or with equivalent concentrations of ethanol solvent. Bacteria were harvested by centrifugation and resuspended twice in PBS to wash cells prior to adjustment to OD_600_=0.1 (∼10^8^ cfu ml^-1^) and bacteria were challenged with varying concentrations of D-sphingosine. Bacteria were sampled at intervals and serially diluted to enumerate viability by culture on agar.

### Sequence data accession

RNA-seq data has been deposited in the ArrayExpress database at EMBL-EBI (www.ebi.ac.uk/arrayexpress) under accession number E-MTAB-9272. Sequence reads of experimentally evolved isolates have been deposited in the European nucleotide archive (ENA) at EMBL-EBI under accession number PRJEB39049.

## Results

### Staphylococcal growth inhibition by D-sphingosine

Sphingosines are potently antimicrobial to staphylococci (Bibel et al., 1992, Fischer et al., 2012), which was confirmed in this study by testing multiple *S. aureus* and *S. epidermidis* strains for their D-sphingosine MIC and MBC (Table 1). Comparison of these data revealed an MIC range from 8-38 μM with MBC values that typically matched, suggesting that once the concentration of D-sphingosine reached a threshold level its activity was lethal. The MIC and MBC values determined here were used to establish conditions for further examining the responses of staphylococci to sphingosines.

**Table 1:** D-sphingosine minimum inhibitory concentrations (MICs) and minimum bactericidal concentrations (MBCs) for *S. aureus* and *S. epidermidis*. Data represent a minimum of 3 independent assays.

| Species | Strain | MIC (μM) | MBC (μM) |
| --- | --- | --- | --- |
| *S. aureus* | SH1000 | 8 | 8 |
|  | Newman | 16 | 38 |
|  | MRSA 252 | 38 | 38 |
|  | MSSA 476 | 38 | 38 |
|  | BL137 | 38 | 38 |
| *S. epidermidis* | Tü3298 | 38 | 38 |
|  | NCTC 1457 | 8 | 8 |
|  | Rp62a | 38 | 38 |
|  | B19 | 8 | 8 |
|  | O16 | 8 | 8 |

### Transcriptomic response to D-sphingosine challenge

A transcriptomics led approach was taken to investigate the outcomes of *S. aureus* and *S. epidermidis* challenge with D-sphingosine and determine whether there were discrete responses. Exponentially growing cultures of *S. epidermidis* Tü3298 and *S. aureus* Newman were challenged with D-sphingosine for 20 min prior to harvesting of cells for RNA-Seq, using an experimental design described previously (Kenny et al., 2009, Moran et al., 2017). A D-sphingosine challenge concentration of 5 μM was chosen as the minimum concentration that the lipid caused growth inhibition of both species under the culture conditions (Figure S1 and data not shown). Controls were treated with the corresponding volume of ethanol solvent, which did not affect growth at the concentration used.

The challenge stimulon of *S. aureus* Newman comprised 1331 genes that were differentially expressed (DE) in response to D-sphingosine, of which 666 were down-regulated and 665 were up-regulated (Fig 1; Table S3). Despite the numerous changes, only 91 genes were DE >log_2_ 2-fold, of which 57 were down-regulated and 34 were up-regulated. Contrastingly, the *S. epidermidis* Tü3298 stimulon showed a more modest number of DE genes in response to D-sphingosine, with only 129 down-regulated and 211 up-regulated genes. Of the DE genes from each species, 1071 had homologs in the other species and in common 101 were up-regulated and 43 were down-regulated (Fig 1, Table S3). Confirmation of DE gene expression from the RNA-Seq was done using qPCR for 5 selected genes (Figure S2).

**Figure 1.**
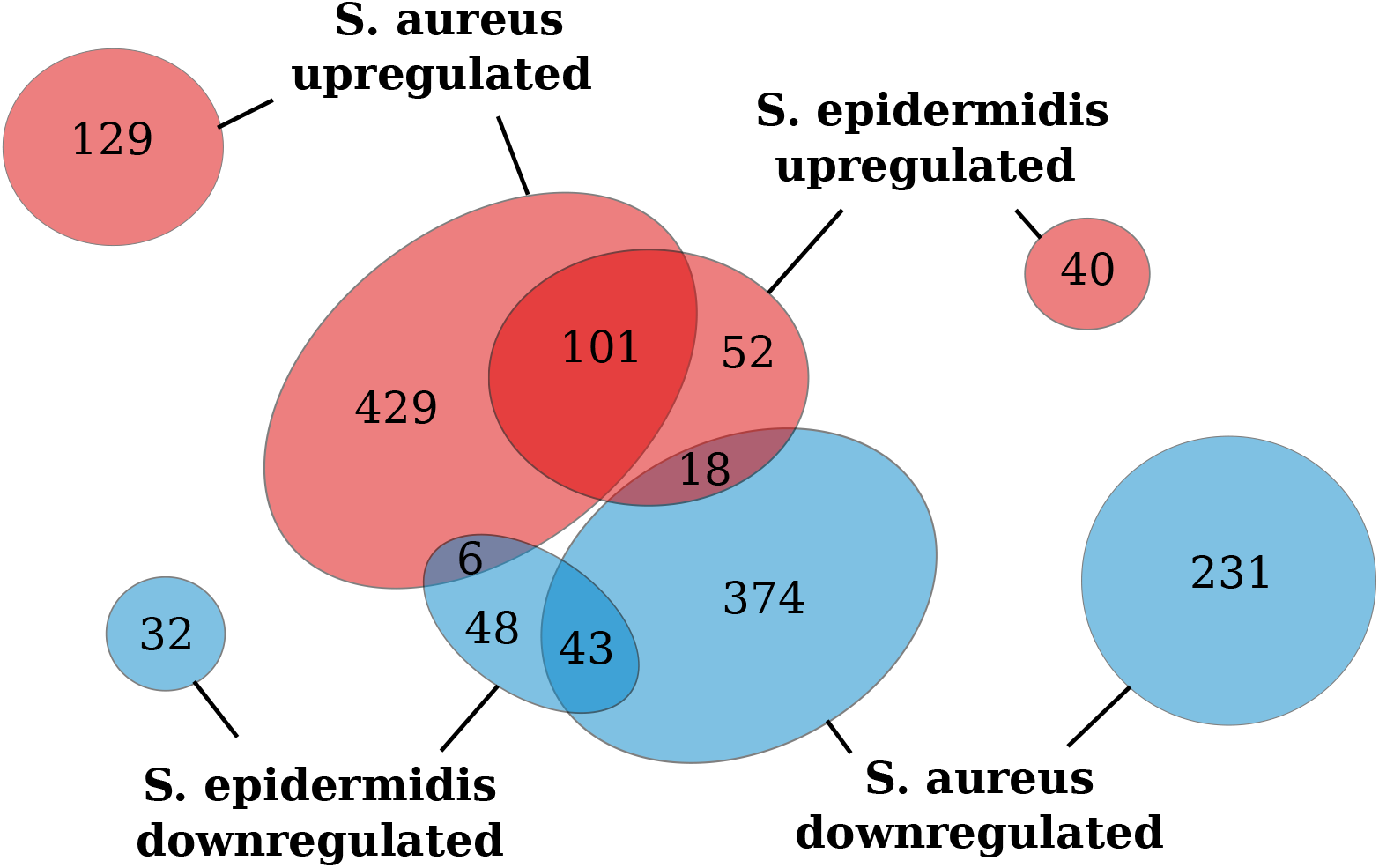
Comparison of up- and down-regulated genes of *S. aureus* and *S. epidermidis* after D-sphingosine challenge. Genes with homologs in both species are indicated in the central ovals, whereas genes without homologues are shown in the peripheral ovals. Common genes are indicated in overlapping portions of the coloured shapes, comprising the D-sphingosine challenge stimulon.

### Phosphate transporter upregulation

Within the common D-sphingosine stimulon of *S. aureus* and *S. epidermidis* (Table S4), the *pstSCAB* phosphate transporter operon was differentially expressed in both species with very high expression in *S. aureus* (log_2_FC 3.1-4.1) (Figure 2). Shared upregulated expression led us to hypothesise that phosphate homeostasis contributes to D-sphingosine resistance or metabolism in staphylococci. The transcription upregulation was confirmed by qPCR for *pstB* of *S. epidermidis* Tü3298 and *S. aureus* Newman, and this upregulation response was conserved across a range of *S. aureus* strains (Figure 2).

**Figure 2.**
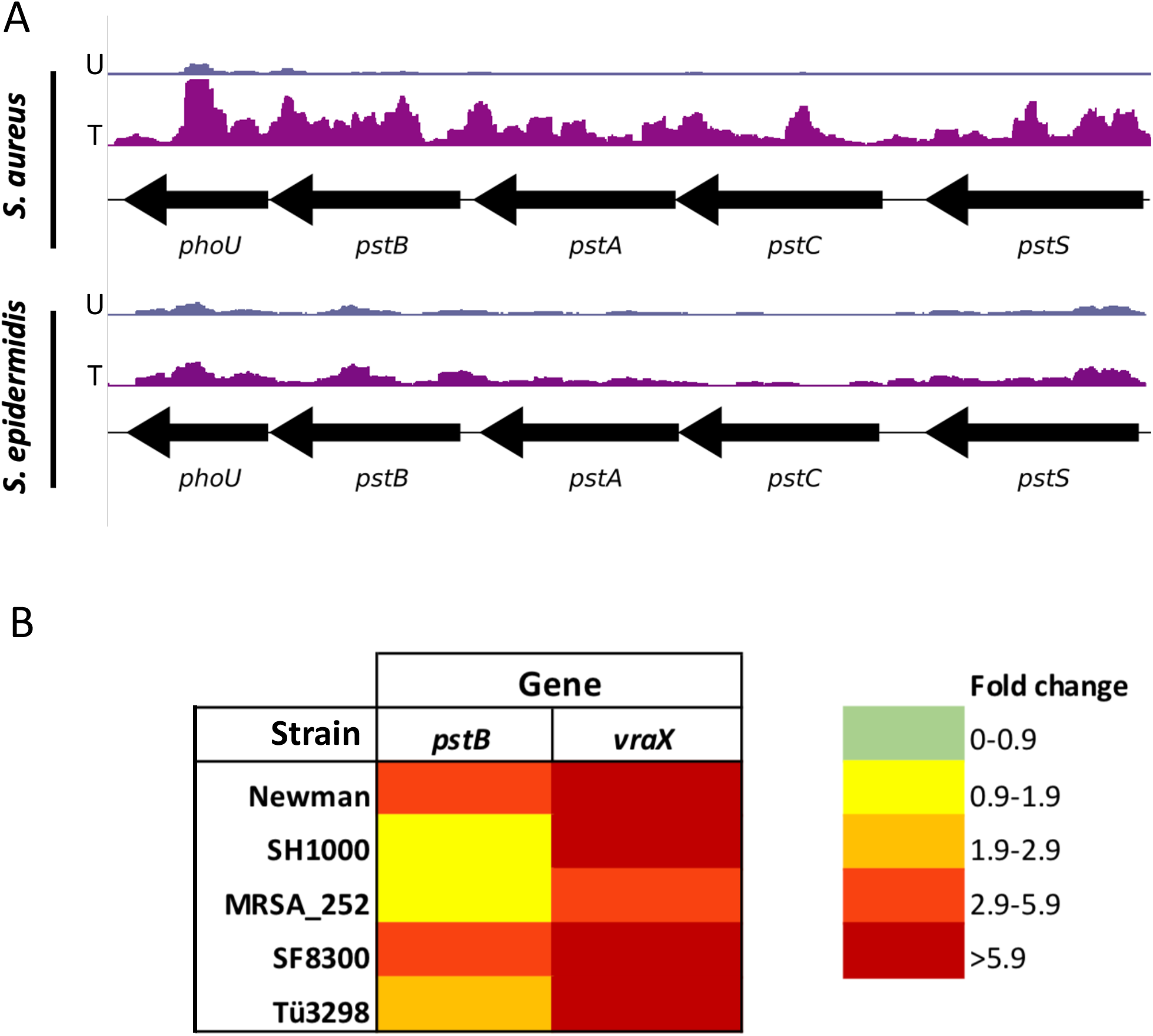
Transcription of key genes post-challenge with sphingosine. (A) Transcription of *pstSCABphoU* for control untreated (U, blue) and treated cells (T, purple) of *S. aureus* and *S. epidermidis*. (B) qPCR of *pstB* and *vraX* across different *S. aureus* and *S. epidermidis* strains showing fold-change after challenge with D-sphingosine.

The contribution of the PstSCAB transport system operon to D-sphingosine survival was tested by constructing an allelic replacement mutant of *pstS* in *S. aureus* Newman. The *pstSCAB::tet* allelic replacement mutant exhibited a lower D-sphingosine MIC relative to its parental strain (6.6 vs 16.5 μM). A *pstS::mariner* transposon mutant exhibited a matching MIC as the operon deletion confirming the phenotype. Additionally, the contribution of downstream operon gene *phoU1* to survival was also tested by generating a strain Newman *phoU1::mariner* transposon mutant, which had a lower MIC compared with its isogenic parent (9.9 μM vs 16.5 μM). Both *phoU1* and *phoU2* are described to contribute to persister formation and virulence gene expression in *S. aureus* and deletion of *phoU2* caused increased phosphate transporter expression (Shang et al., 2020).

A *pstSCAB* operon complementation strain could not be generated in *S. aureus* despite multiple strategies. Presumably the membrane protein expression of the operon in a multicopy plasmid was toxic in *E. coli*. We have observed this phenomenon previously with *S. aureus* transporter proteins (Horsburgh et al., 2004, Kenny et al., 2013) and we were unable to circumvent this issue by cloning the *pstSCAB* operon directly into *S. aureus* RN4220, a strategy that we employed previously.

### Altered TCS and virulence gene expression

Sphingosine challenge led to downregulation of the SaeSR two-component system (TCS) genes *saeS* and *saeR* and upregulation of the VraSR TCS genes *vraS* and *vraR* of both *S. aureus* Newman and *S. epidermidis* Tü3298. In *S. aureus*, this was accompanied by downregulation of multiple virulence factor genes encoding adhesins, immunomodulatory factors and toxins (Table S3). Many of these factors promote host colonisation as well as virulence and are part of the SaeSR regulon (Geiger et al., 2008, Liu et al., 2016) that could suggest D-sphingosine alters TCS activation. However, *S. aureus* Newman has a point mutation in *saeS* that results in constitutive activation of the Sae system regulator SaeR (Cue et al., 2015), reducing clarity that the SaeSR regulon downregulation is a result of D-sphingosine interaction. Increased expression of *vraSR* was observed together with its cell wall stress stimulon (e.g. *pbpB, murZ*, and *msrA1*; Table S3). The capacity of VraSR to regulate the response of *S. aureus* to D-sphingosine was supported with the reduced MIC of both *vraS::tet* (6.6 mM) and *vraR::tet* (6.6 mM) allelic replacement mutants compared with their isogenic parent SH1000 (16.3 μM).

In *S. epidermidis, sepA* (SETU_02085), encoding a metalloprotease that inhibits LL-37 activity (Paharik et al., 2017), was the only DE virulence-associated gene other than *saeS* and *saeR*. While SaeRS TCS enhances virulence in *S. epidermidis*, its role is primarily linked to anaerobic growth and control of autolysis rather than expression of virulence determinants (Handke et al., 2008, Lou et al., 2011).

### Cell wall and membrane gene expression

There was a pronounced difference between *S. epidermidis* and *S. aureus* gene expression for glycerolipid and fatty acid biosynthesis and degradation pathways (Table 2; S3). In response to D-sphingosine challenge of *S. aureus*, expression of fatty acid biosynthesis and degradation pathway genes and glycerophospholipid biosynthesis genes were mostly upregulated. Contrastingly, there was little altered gene expression in these pathways with *S. epidermidis*. Many *S. aureus* peptidoglycan biosynthesis genes were upregulated in contrast to *S. epidermidis* (Table 2; S3). Increased cell wall and cell membrane biosynthesis is typically associated with cell growth, although the culture data indicated a minor impact following D-sphingosine challenge suggesting instead there could be restructuring of the cell wall and membrane in response to the imposed or consequent stress. Upregulation of the VraSR cell wall stress TCS indicates a potential mechanism for DE of *S. aureus* cell wall stress regulon genes. The stress impact of D-sphingosine could be evidenced in both staphylococci by upregulation of the gene for the stress response regulator CtsR, and its regulon genes *dnaJ* and *dnaK* (Table S3, S4).

**Table 2.** Differential gene expression of lipid and fatty acid catabolism and metabolism genes after D-sphingosine challenge. Colouring indicates the extent of log_2_ fold DE, with yellow to red indicating low to high upregulation and pale blue to dark blue indicating low to high downregulation.

| *S. aureus* |  | *S. epidermidis* |  |
| --- | --- | --- | --- |
| LogFC | Gene name |  | LogFC |
| Glycerolipid biosynthesis |  |  |  |
| -0.6 | *plsK* |  |  |
| -0.6 | *plsX* |  |  |
| 0.9 | *glpK* |  |  |
| -2.6 | *geh* | *SETU_02128* | 1.9 |
| -1.8 | *lip* |  |  |
| Glycerol, glycerolipid, glycerophospholipid metabolism |  |  |  |
| 0.9 | *NWMN_1858* | *SETU_01602* |  |
| -0.6 | *plsK* |  |  |
| -0.6 | *plsX* |  |  |
| 1.3 | *glpQ* |  |  |
| 0.8 | *NWMN_0984* |  |  |
| 1 | *NWMN_1614* |  |  |
| Glycerol, glycerolipid, glycerophospholipid metabolism |  |  |  |
| 1.7 | *NWMN_2331* | *SETU_01869* | 1 |
| 0.5 | *NWMN_0711* |  |  |
| 0.7 | *NWMN_0619* | *SETU_02171* | 1 |
| 0.6 | *NWMN_0620* | *SETU_02170* | 1.3 |
| 1.6 | *glpD* |  |  |
| 0.5 | *NWMN_1230* |  |  |
| -0.6 | *NWMN_1926* |  |  |
| Fatty acid degradation |  |  |  |
| 0.9 | *NWMN_1858* | *SETU_01602* |  |
| 1 | *fadE* |  |  |
| 0.8 | *NWMN_0346* | *SETU_00013* |  |
| -0.7 | *adhE* |  |  |
| -2.3 | *adh1* | *SETU_00229* | 1.6 |
| 0.9 | *NWMN_2091* | *SETU_01662* | -0.8 |
| 0.6 | *fadA* |  |  |
| 0.6 | *fadB* |  |  |
| 0.9 | *accC* |  |  |
| 0.8 | *accB* |  |  |
| -0.5 | *fadD* |  |  |
| -1 | *NWMN_0853* |  |  |
| -0.8 | *NWMN_0854* |  |  |
| -0.5 | *fabG* |  |  |

With respect to altering cell surface features, both *S. aureus* and *S. epidermidis* responded to D-sphingosine with DE of cell membrane and cell wall modification genes. In *S. epidermidis* this included down-regulation of *mprF* and the *dlt* operon, which encode enzymes that lysinylate peptidoglycan and D-alanylate teichoic acids respectively (Weidenmaier et al., 2005), which could potentially lead to an overall increase of negative surface charge. The *S. aureus* response reveals up-regulated genes of the *tag* operon, which encodes biosynthesis of wall teichoic acid and its attachment. The upregulation of this modification could similarly increase negative cellular charge, particularly if combined with cell wall thickening caused by increased peptidoglycan biosynthesis. At this stage, it is unclear how the amphipathic nature of the sphingosine combined with its positively charged amino group (at neutral pH) relates to these changes.

*S. aureus* downregulated expression of carotenoid biosynthesis (*crt*MNOPQ) genes between 1.15-1.74 log_2_ fold after sphingosine challenge. Modulated pigment expression is a known response to membrane and peroxide stress (Horsburgh et al., 2002, Liu et al., 2005). Carotenoid levels were too low to detect variations at the mid-log growth phase used for RNA-Seq expression data, however after 24 h of growth in the presence of 5 μM D-sphingosine there was increased carotenoid production in both tested *S. aureus* strains (data not shown). Consistent with increased carotenoid after overnight culture with sphingosine, cells had a greater survival from 1 mM hydrogen peroxide challenge (data not shown).

### Evolution of resistance to D-sphingosine

While transcriptomic data can be used to identify resistance responses it has limitations and we supplemented our investigation with an experimental evolution approach to select for *S. aureus* and *S. epidermidis* mutants with increased resistance to D-sphingosine.

*S. aureus* SH1000 and *S. epidermidis* Rp62a were growth-passaged in increasing concentrations of D-sphingosine beyond the day when maximum MIC values were reached. Both species developed stable phenotypes over time that resulted in increased growth in elevated D-sphingosine (Fig 3). For undetermined reasons the maximum sphingosine concentration at which *S. aureus* cultures could grow was markedly higher and was reached more rapidly. The MIC values of *S. aureus* isolates from days 3 and 5 showed a 2 to16-fold MIC increase (Fig 3) and although bacteria were passaged for 9 days, there was no additional increase in MIC beyond day 5, indicating a maximum threshold for the conditions tested.

**Figure 3.**
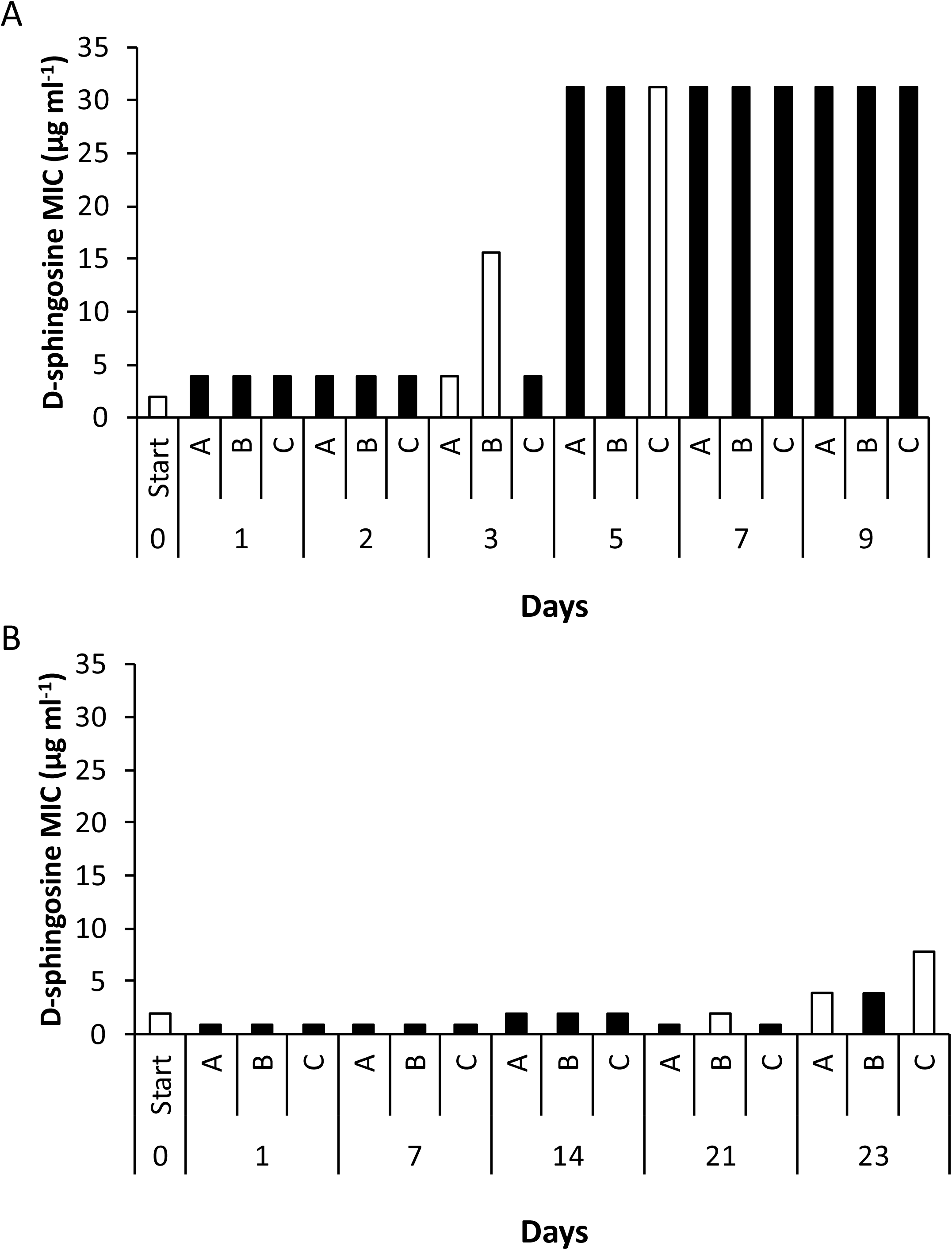
D-sphingosine MIC of experimentally evolved isolates. *S. aureus* (A) and *S. epidermidis* (B) from passage days 0-23. For each day of sampling, individual isolates of separate growth passages are labelled A, B, C. Grey bars indicate the representative isolates selected for whole genome sequencing.

*S. epidermidis* cultures could grow with increased concentrations of D-sphingosine from day 2 when serially passaged (data not shown), but this was not matched with increased MIC when testing evolved isolates. This growth was likely due to adaptive changes or unstable genetic mutations that quickly reverted without the selective pressure of D-sphingosine. By day 23, increased MIC of isolates became fixed with 2 to 4-fold MIC increases. Whole genome sequencing was used to identify mutations contributing to the phenotype of representative isolates of *S. aureus* with 2, 8 and 16-fold increased MIC, and *S. epidermidis* isolates with 2 and 4-fold increased MIC. An *S. epidermidis* evolved isolate with the same MIC as WT was selected as an internal control, to resolve mutations that did not contribute to increased MIC.

### Genetic variation in evolved isolates

Genome sequencing of *S. aureus* and *S. epidermidis* isolates with experimentally evolved resistance to D-sphingosine identified single nucleotide polymorphisms (SNPs) (Fig 3; 4; Table S5). Most mutations were present in more than one isolate. Common to both species with increased D-sphingosine MIC were SNPs in *farE* and/or *farR* (Fig 4). In addition, the isolate with the highest D-sphingosine MIC, *S. aureus* SH1000-5C, had a SNP in *farE* (FarE-V296A) not shared by any other sequenced isolate (Table S5). FarR is a TetR-like regulator of the efflux pump FarE, known to be involved in resistance to another antimicrobial skin lipid, linoleic acid (Kenny et al., 2009, Alnaseri et al., 2015). The FarE/FarR regulator-efflux system was therefore considered a strong candidate resistance mechanism to D-sphingosine and was investigated further for its role in sphingosine resistance.

**Figure 4.**
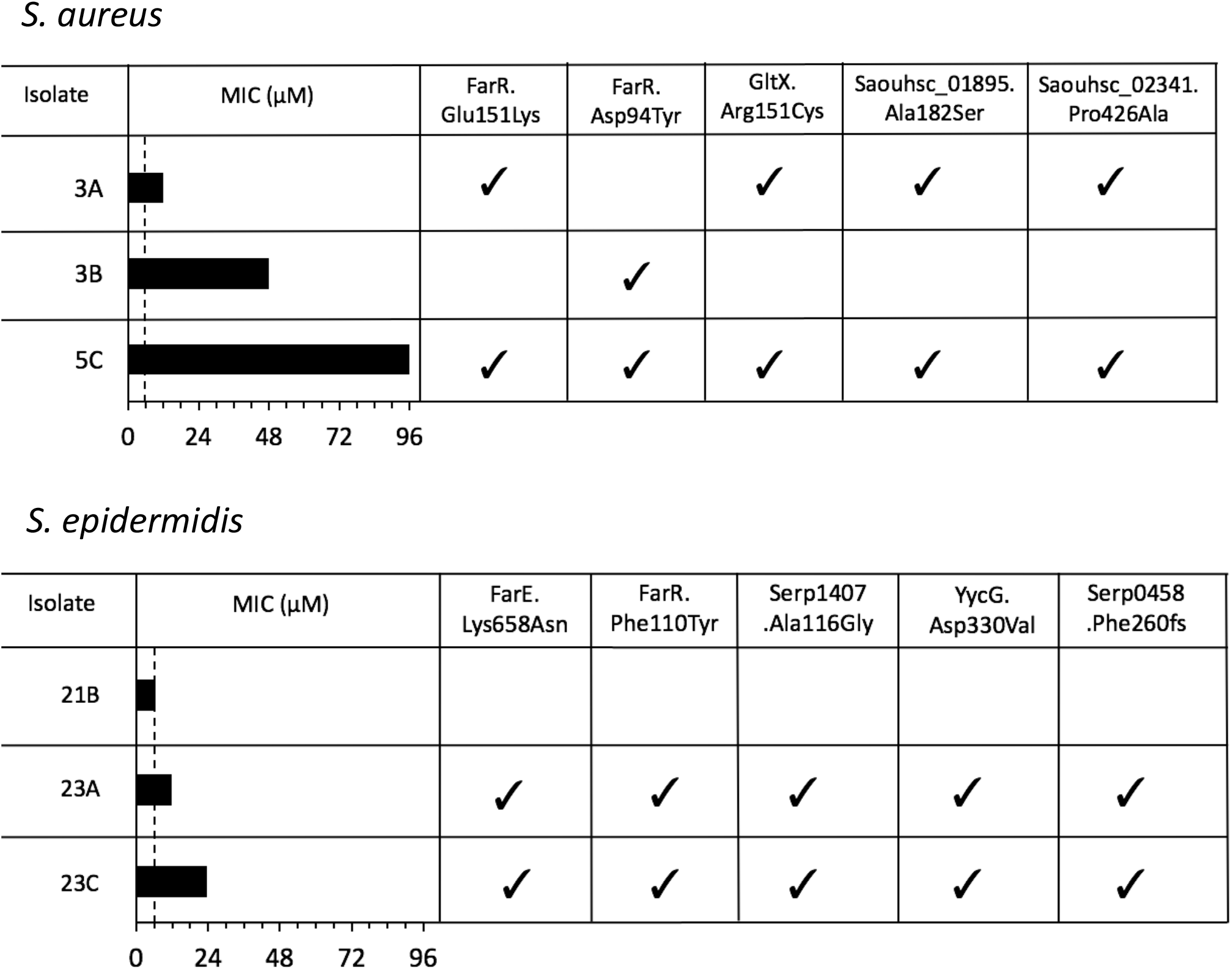
Variants and MICs of *S. aureus and S. epidermidis* experimental evolution isolates. Variants that occurred in more than one isolate, within a gene and caused non-synonymous changes are shown. Dotted line indicates D-sphingosine MIC of WT

### The contribution of FarE/R in D-sphingosine Resistance

To investigate whether both *S. aureus* FarE and FarR contribute to D-sphingosine resistance, a *farR* mutant was constructed and investigated alongside a previously constructed *farE* mutant (Kenny et al., 2009). Overnight culture of *S. aureus* with 5 μM D-sphingosine led to adaptations that enhanced survival; untreated *S. aureus* challenged with 15 μM D-sphingosine had 70% survival at 15 min, whereas D-sphingosine precultured *S. aureus* had 100% survival at 15 min. In contrast, *farE* and *farR* mutants showed 0% survival with challenge at 15 μM lipid with or without preculture.

Compared with wild-type parent strain MIC (16.3 μM), gene deletion caused reduced MIC: *farE* (9.9 μM) and *farR* (11.8 μM). To resolve whether the SNPs identified in the experimental evolution in *farR* conferred increased resistance to D-sphingosine, the *farR*-D94Y-E151K variant gene from SH1000-5C was introduced into SH1000 WT. The verified mutant, SH1000 FarR-D94Y-E151K, exhibited a 10-fold increase in MIC (165 μM).

## Discussion

*Staphylococcus* species are known for their particular niche success in colonisation of skin, an ability which is multifactorial. The various lipids and fatty acids that are produced from epidermal and sebaceous compartments form a key barrier to colonisation of skin. Study here of the staphylococcal response to D-sphingosine was made to gain insight of the mechanisms that could overcome activity of this ceramide breakdown product.

Using an RNA-Seq approach, it was determined that both *S. aureus* and *S. epidermidis* responded to D-sphingosine challenge using several homologous response networks. Upregulation of VraSR and CtsR and downregulation of SaeSR regulons appear within these data reflecting a stress stimulus response to cell surface challenge. Similar to the effects of linoleic and sapienic acids that affect SaeSR regulation (Kenny et al., 2013, Moran et al., 2017), D-sphingosine may contribute to modulating *S. aureus* colonisation factors.

Upregulation of both VraSR and CtsR regulons could map to the mode of action of sphingosines. Sphingosine was recently shown to interact with the cell membrane via cardiolipin to kill *S. aureus* through rapid membrane permeabilization (Verhaegh et al., 2020). The authors speculate this could be caused by the cardiolipin-sphingosine interaction forming rigid domains within the membrane, as is indicated for the aminoglycoside 3’, 6-dinonylneamine (Verhaegh et al., 2020, El Khoury et al., 2017). Both *S. aureus* and *S. epidermidis* respond to D-sphingosine by modulating lipid biosynthesis and fatty acid degradation pathways in dissimilar expression patterns, but there was no alteration to cardiolipin biosynthesis gene transcription, with neither *cls1* nor *cls2* genes being DE.

A common response to sphingosine in both staphylococci was upregulation of the transport operon *pstSCAB*. The encoded phosphate-specific transport system is a high-affinity, low-velocity, free-Pi transporter system that is structurally similar to ABC transporters (Ames, 1986). Downstream of the *pstSCAB* operon, the *phoU1* gene encodes a transporter-associated protein. PhoU proteins contribute to multidrug tolerance and persistence in *S. aureus* and impacts on widespread processes beyond inorganic phosphate (Pi) transport (Li and Zhang, 2007). Upregulation of proteins encoded by the *pstSCAB* and *phoU* operons were also found in antimicrobial peptide ranalexin resistance in methicillin-resistant *S. aureus* (Overton et al., 2011). Genetic disruption of the Pi transporter, PstC, had no effect on ranalexin sensitivity, suggesting that MRSA adopts a PhoU-mediated persister phenotype to acquire antimicrobial tolerance, and upregulation of the Pi transporter is not a major component of this response (Overton et al., 2011). However, inactivation of *phoU2* did lead to upregulation of phosphate transport operon expression (Shang et al., 2020). Inactivation of the *pstSCAB* operon in *S. aureus* was shown here to reduce the sphingosine MIC, upregulated phosphate transport could allow staphylococci to incorporate sphingosine into phospholipid biosynthesis, but this was not tested or explored further.

To further identify mechanisms of survival from sphingosine, we employed experimental evolution in the presence of sphingosine. Genome sequencing of isolates with greater MIC to sphingosine revealed patterns of SNPs common to both *S. aureus* and *S. epidermidis* that highlighted the FarE-FarR transporter-regulator proteins could contribute to survival. *S. aureus* FarE was previously identified as a lipid transporter that provides enhanced tolerance to the antimicrobial lipid linoleic acid (Kenny et al., 2009, Alnaseri et al., 2015). *S. aureus* FarR regulates FarE in *S. aureus* (Alnaseri et al., 2015, Nguyen et al., 2019) and the *S. epidermidis* gene for the FarR homologue SERP0247 and efflux transporter gene SERP0245 share synteny. Inactivation of *farE* or *farR* in *S. aureus* reduced the D-sphingosine MIC, and introduction of the double SNP mutations *farR*-D94Y-E151K into *S. aureus* SH1000 increased MIC 10-fold to 165 μM. Examination of the sphingosine challenge transcriptome shows only the lipid A and amphipathic drug ABC transporter MsbA (Velamakanni et al., 2008) as a potential exclusion transport mechanism that was upregulated; its role was not tested in the present study and was not identified in the experimental evolution approach.

FarR is a member of the TetR family of repressors that form homodimers. Binding of the substrate alters the structure of TetR proteins, reducing their affinity for the DNA binding site and relieving repression of the target gene (Ramos et al., 2005). Most of the *S. aureus* and *S. epidermidis* FarR mutations are predicted to be located at the entrance to and within a pocket of the substrate-binding domain. Therefore, the identified changes could either affect how this repressor interacts with its substrate or cause the protein to be in the conformation with low DNA affinity independent of the substrate. A further *S. aureus* mutation, yielding FarR.A40S, occurred in the α-2 region of the DNA binding domain that may alter DNA binding. The transcriptional effects of a C116R *farR* mutation were identified in a study of experimentally evolved resistance to rhodomyrtone, a plant acylphloroglucinol antimicrobial, with upregulated FarE expression as the resistance mechanism (Nguyen et al., 2019). Of note, this study revealed that the selected mutant had greater virulence in murine models, which demands further study of sphingosine selected mutants.

The FarE transporter was not a regulated exclusion mechanism in response to sphingosine and the mutations require future investigation to identify their exact outcomes. Cellular resistance gained through SNPs in the *farE* efflux transporter gene could alter affinity (K_m_) for D-sphingosine and/or flux (V_max_). The FarE transporter is a member of the RND protein family that encode secondary multidrug transporters for a wide range of compounds including drugs, metals, solvents, fatty acids, detergents and dyes (Domenech et al., 2005, Putman et al., 2000). The described RND proteins are Mmpl-like transporters and some are described to act as “membrane hoovers” that efflux toxic substrates from the membrane (Truong-Bolduc et al., 2014). Transmembrane domains of the *E. coli* multimer AcrB appear to interact loosely, creating vestibules allowing phospholipids and other membrane components access to the central pore which is used for substrate transportation (Murakami et al., 2002). The creation of vestibules through loose interactions of transmembrane domains could explain how substrates in the membrane enter the pump. RND family proteins are characterised by their structure of 12 transmembrane domains, with the 1^st^ and 2^nd^, 7^th^ and 8^th^ being linked by extracytoplasmic loops, which likely confer substrate specificity (Domenech et al., 2005, Varela et al., 2012).

Both mutant staphylococcal FarE proteins (SERP0245 and SAOUHSC_02866) in this study would bear a single amino acid change facing outwards in an α-helix within the portion of the 12th and 6^th^ transmembrane domains (respectively) nearest to the extracellular domains. These domains are proposed to have a conserved sequence (Putman et al., 2000), and may contribute to substrate interaction. Indeed, Guay *et al*. (1994) identified amino acid changes in transmembrane helices of the RND transporter *tetA*(B) that increased resistance to 9-(dimethylglycylamido)minocycline whilst decreasing tetracycline resistance. This change in resistance was likely due to differences in substrate specificity (Guay et al., 1994) and was predicted to occur at similar depth within the membrane to the changes observed here, though in different helices.

Human skin colonisation by bacterial genera requires investigation to identify their evolved niche specialisations. Such studies will unlock the interplay with the host and reveal balances between commensalism and pathogenesis that could also identify causes of microbiome population changes described as dysbiosis. Concluding from this study of sphingosine survival with these two clinically-important staphylococci, the administration of sphingosines to ameliorate topical skin diseases could result in high-level resistance through selection of mutations in genes encoding the FarE-FarR lipid efflux system.

## Acknowledgements

YC was funded with support from a University of Liverpool bursary. JCM was funded by BBSRC grants BB/D003563/1 and BB/L023040/1 awarded to MJH, both with support from Unilever Plc. The funders were not involved in the study design, collection of samples, analysis of data, interpretation of data, the writing of this report or the decision to submit this report for publication. The Centre for Genomic Research, University of Liverpool, UK carried out rRNA-Seq data generation and processing.

---

**Figure S1. D-sphingosine growth effects on *S. aureus*.** Bacteria cultures in THB medium were challenged with D-sphingosine at 5-7 μM during growth at OD600=0.5. Control was ethanol solvent only (0.3 mM) and blank was unamended bacteria.

**Figure S2. Relative DE for selected genes of *S. aureus* after challenge with D-sphingosine.** Black bars indicate the fold change from RNA-Seq data, grey bars indicate fold change from qPCR validation. Error bars refer to standard deviation of fold changes of three replicates.

**Table S1. Primers used in study.**

**Table S2. Sequence mapping data of experimentally evolved isolates of S. aureus and S. epidermidis.**

**Table S3. DE genes of *S. aureus* and *S. epidermidis* in response to D-sphingosine.** Genes with differential expression in RNA Seq analysis between control and D-sphingosine are shown for each species, as well as comparison of all homologous genes between species.

**Table S4. Common D-sphingosine stimulon of *S. aureus* and *S. epidermidis*.** Common DE homologous genes with the same trend of regulation in response to D-sphingosine challenge of *S. aureus* Newman and *S. epidermidis* Tü3298. DE genes are listed and values are shaded using a heatmap as per the indicated key.

**Table S5. Mutations in sequenced D-sphingosine evolved *S. aureus* and *S. epidermidis*.** All mutations found in evolved isolates not found in parental genomes.

## References

Alnaseri, H., Arsic, B., Schneider, J. E., Kaiser, J. C., Scinocca, Z. C., Heinrichs, D. E. & Mcgavin, M. J. 2015. Inducible expression of a Resistance-Nodulation-Division-type efflux pump in *Staphylococcus aureus* provides resistance to linoleic and arachidonic acids. J Bacteriol, 197, 1893–905.

Ames, G. 1986. Bacterial periplasmic transport systems: structure, mechanism, and evolution. Ann Rev Biochem, 55, 397–425.

Arikawa, J., Ishibashi, M., Kawashima, M., Takagi, Y., Ichikawa, Y. & Imokawa, G. 2002. Decreased levels of sphingosine, a natural antimicrobial agent, may be associated with vulnerability of the stratum corneum from patients with atopic dermatitis to colonization by *Staphylococcus aureus*. J Invest Dermatol, 119, 433–9.

Beck, S., Sehl, C., Voortmann, S., Verhasselt, H. L., Edwards, M. J., Buer, J., Hasenberg, M., Gulbins, E. & Becker, K. A. 2020. Sphingosine is able to prevent and eliminate *Staphylococcus epidermidis* biofilm formation on different orthopedic implant materials in vitro. J Mol Med (Berl), 98, 209–19.

Bibel, D. J., Aly, R. & Shinefield, H. R. 1992. Antimicrobial activity of sphingosines. J Invest Dermatol, 98, 269–73.

Coates, R., Moran, J. & Horsburgh, M. J. 2014. Staphylococci: colonizers and pathogens of human skin. Future Microbiol, 9, 75–91.

Coates-Brown, R., Moran, J. C., Pongchaikul, P., Darby, A. C. & Horsburgh, M. J. 2018. Comparative genomics of Staphylococcus reveals determinants of speciation and diversification of antimicrobial defense. Front Microbiol, 9, 2753.

Cue, D., Junecko, J. M., Lei, M. G., Blevins, J. S., Smeltzer, M. S. & Lee, C. Y. 2015. SaeRS-dependent inhibition of biofilm formation in *Staphylococcus aureus* Newman. PLoS One, 10, e0123027.

Domenech, P., Reed, M. B. & Barry, C. E., 3RD 2005. Contribution of the *Mycobacterium tuberculosis* MmpL protein family to virulence and drug resistance. Infect Immun, 73, 3492–501.

El Khoury, M., Swain, J., Sautrey, G., Zimmermann, L., Van der Smissen, P., Decout, J. L. & Mingeot-Leclercq, M. P. 2017. Targeting bacterial cardiolipin enriched microdomains: an antimicrobial strategy used by amphiphilic aminoglycoside antibiotics. Sci Rep, 7, 10697.

Fey, P. D., Endres, J. L., Yajjala, V. K., Widhelm, T. J., Boissy, R. J., Bose, J. L. & Bayles, K. W. 2013. A genetic resource for rapid and comprehensive phenotype screening of nonessential *Staphylococcus aureus* genes. mBio, 4, e00537–12.

Fischer, C. L., Drake, D. R., Dawson, D. V., Blanchette, D. R., Brogden, K. A. & Wertz, P. W. 2012. Antibacterial activity of sphingoid bases and fatty acids against Gram-positive and Gram-negative bacteria. Antimicrob Agents Chemother, 56, 1157–61.

Geiger, T., Goerke, C., Mainiero, M., Kraus, D. & Wolz, C. 2008. The virulence regulator Sae of *Staphylococcus aureus:* promoter activities and response to phagocytosis-related signals. J Bacteriol, 190, 3419–28.

Guay, G., Tuckman, M. & Rothstein, D. 1994. Mutations in the *tetA*(B) gene that cause a change in substrate specificity of the tetracycline efflux pump. Antimicrob Agents Chemother, 38, 857–60.

Handke, L. D., Rogers, K. L., Olson, M. E., Somerville, G. A., Jerrells, T. J., Rupp, M. E., Dunman, P. M. & Fey, P. D. 2008. *Staphylococcus epidermidis saeR* is an effector of anaerobic growth and a mediator of acute inflammation. Infect Immun, 76, 141–52.

Horsburgh, M. J., Ingham, E. & Foster, S. J. 2001. In *Staphylococcus aureus, fur* is an interactive regulator with PerR, contributes to virulence, and Is necessary for oxidative stress resistance through positive regulation of catalase and iron homeostasis. J Bacteriol, 183, 468–75.

Horsburgh, M. J., Wharton, S. J., Cox, A. G., Ingham, E., Peacock, S. & Foster, S. J. 2002. MntR modulates expression of the PerR regulon and superoxide resistance in *Staphylococcus aureus* through control of manganese uptake. Mol Microbiol, 44, 1269–86.

Horsburgh, M. J., Wiltshire, M. D., Crossley, H., Ingham, E. & Foster, S. J. 2004. PheP, a putative amino acid permease of *Staphylococcus aureus,* contributes to survival *in vivo* and during starvation. Infect Immun, 72, 3073–6.

Kenny, J. G., Moran, J., Kolar, S. L., Ulanov, A., Li, Z., Shaw, L. N., Josefsson, E. & Horsburgh, M. J. 2013. Mannitol utilisation is required for protection of *Staphylococcus aureus* from human skin antimicrobial fatty acids. PLoS One, 8, e67698.

Kenny, J. G., Ward, D., Josefsson, E., Jonsson, I. M., Hinds, J., Rees, H. H., Lindsay, J. A., Tarkowski, A. & Horsburgh, M. J. 2009. The *Staphylococcus aureus* response to unsaturated long chain free fatty acids: survival mechanisms and virulence implications. PLoS One, 4, e4344.

Langmead, B. & Salzberg, S. L. 2012. Fast gapped-read alignment with Bowtie 2. Nat Methods, 9, 357–9.

Li, Y. & Zhang, Y. 2007. PhoU is a persistence switch involved in persister formation and tolerance to multiple antibiotics and stresses in *Escherichia coli*. Antimicrob Agents Chemother, 51, 2092–9.

Liu, G. Y., Essex, A., Buchanan, J. T., Datta, V., Hoffman, H. M., Bastian, J. F., Fierer, J. & Nizet, V. 2005. *Staphylococcus aureus* golden pigment impairs neutrophil killing and promotes virulence through its antioxidant activity. J Exp Med, 202, 209–15.

Liu, Q., Yeo, W. S. & Bae, T. 2016. The SaeRS Two-Component System of *Staphylococcus aureus*. Genes, 7.

Lou, Q., Zhu, T., Hu, J., Ben, H., Yang, J., Yu, F., Liu, J., Wu, Y., Fischer, A., Francois, P., Schrenzel, J. & Qu, D. 2011. Role of the SaeRS two-component regulatory system in *Staphylococcus epidermidis* autolysis and biofilm formation. BMC Microbiol, 11, 146.

Mccarthy, D. J., Chen, Y. & Smyth, G. K. 2012. Differential expression analysis of multifactor RNA-Seq experiments with respect to biological variation. Nucleic Acids Res, 40, 4288–97.

Moran, J. C., Alorabi, J. A. & Horsburgh, M. J. 2017. Comparative transcriptomics reveals discrete survival responses of *S. aureus* and *S. epidermidis* to sapienic acid. Front Microbiol, 8.

Murakami, M., Ohtake, T., Dorschner, R. A., Schittek, B., Garbe, C. & Gallo, R. L. 2002. Cathelicidin anti-microbial peptide expression in sweat, an innate defense system for the skin. J Invest Dermatol, 119, 1090–5.

Nguyen, M. T., Saising, J., Tribelli, P. M., Nega, M., Diene, S. M., Francois, P., Schrenzel, J., Sproer, C., Bunk, B., Ebner, P., Hertlein, T., Kumari, N., Hartner, T., Wistuba, D., Voravuthikunchai, S. P., Mader, U., Ohlsen, K. & Gotz, F. 2019. Inactivation of*farR* causes high rhodomyrtone resistance and increased pathogenicity in *Staphylococcus aureus*. Front Microbiol, 10, 1157.

Nicolaides, N. 1974. Skin lipids: their biochemical uniqueness. Science, 186, 19–26.

Overton, I., Graham, S., Gould, K., Hinds, J., Botting, C., Shirran, S., Barton, G. & Coote, P. 2011. Global network analysis of drug tolerance, mode of action and virulence in methicillin-resistant *S. aureus*. BMC Syst Biol, 5.

Paharik, A. E., Kotasinska, M., Both, A., Hoang, T. N., Buttner, H., Roy, P., Fey, P. D., Horswill, A. R. & Rohde, H. 2017. The metalloprotease SepA governs processing of accumulation-associated protein and shapes intercellular adhesive surface properties in *Staphylococcus epidermidis*. Mol Microbiol, 103, 860–874.

Pappas, A. 2009. Epidermal surface lipids. Dermatoendocrinol, 1, 72–6.

Parsons, J. B., Yao, J., Frank, M. W., Jackson, P. & Rock, C. O. 2012. Membrane disruption by antimicrobial fatty acids releases low-molecular-weight proteins from *Staphylococcus aureus*. J Bacteriol, 194, 5294–304.

Pewzner-JUNG, Y., Tavakoli TABAZAVAREH, S., Grassme, H., Becker, K. A., Japtok, L., Steinmann, J., Joseph, T., Lang, S., Tuemmler, B., Schuchman, E. H., Lentsch, A. B., Kleuser, B., Edwards, M. J., Futerman, A. H. & Gulbins, E. 2014. Sphingoid long chain bases prevent lung infection by *Pseudomonas aeruginosa*. EMBO Mol Med, 6, 1205–14.

Putman, M., Van Veen, H. & Konings, W. 2000. Molecular properties of bacterial multidrug transporters. Microbiol Mol Biol Rev, 64, 672–93.

Ramos, J. L., Martinez-Bueno, M., Molina-Henares, A. J., Teran, W., Watanabe, K., Zhang, X., Gallegos, M. T., Brennan, R. & Tobes, R. 2005. The TetR family of transcriptional repressors. Microbiol Mol Biol Rev, 69, 326–56.

Robinson, M. D., Mccarthy, D. J. & Smyth, G. K. 2010. edgeR: a Bioconductor package for differential expression analysis of digital gene expression data. Bioinformatics, 26, 139–40.

Seitz, A. P., Schumacher, F., Baker, J., Soddemann, M., Wilker, B., Caldwell, C. C., Gobble, R. M., Kamler, M., Becker, K. A., Beck, S., Kleuser, B., Edwards, M. J. & Gulbins, E. 2019. Sphingosine-coating of plastic surfaces prevents ventilator-associated pneumonia. J Mol Med, 97, 1195–1211.

Shang, Y., Wang, X., Chen, Z., Lyu, Z., Lin, Z., Zheng, J., Wu, Y., Deng, Q., Yu, Z., Zhang, Y. & Qu, D. 2020. *Staphylococcus aureus* PhoU homologs regulate persister formation and virulence. Front Microbiol, 11, 865.

Tavakoli Tabazavareh, S., Seitz, A., Jernigan, P., Sehl, C., Keitsch, S., Lang, S., Kahl, B. C., Edwards, M., Grassme, H., Gulbins, E. & Becker, K. A. 2016. Lack of sphingosine causes susceptibility to pulmonary *Staphylococcus aureus* infections in Cystic Fibrosis. Cell Physiol Biochem, 38, 2094–102.

Truong-Bolduc, Q. C., Villet, R. A., Estabrooks, Z. A. & Hooper, D. C. 2014. Native efflux pumps contribute resistance to antimicrobials of skin and the ability of *Staphylococcus aureus* to colonize skin. J Infect Dis, 209, 1485–93.

Varela, C., Rittmann, D., Singh, A., Krumbach, K., Bhatt, K., Eggeling, L., Besra, G. S. & Bhatt, A. 2012. MmpL genes are associated with mycolic acid metabolism in mycobacteria and corynebacteria. Chem Biol, 19, 498–506.

Velamakanni, S., Yao, Y., Gutmann, D. & Van Veen, H. 2008. Multidrug transport by the ABC transporter Sav1866 from *Staphylococcus aureus*. Biochem, 47, 9300–8.

Verhaegh, R., Becker, K. A., Edwards, M. J. & Gulbins, E. 2020. Sphingosine kills bacteria by binding to cardiolipin. J Biol Chem, 295, 7686–7696.

Weidenmaier, C., Peschel, A., Xiong, Y., Kristian, S., Dietz, K., Yeaman, M. & Bayer, A. 2005. Lack of wall teichoic acids in *Staphylococcus aureus* leads to reduced interactions with endothelial cells and to attenuated virulence in a rabbit model of endocarditis. J Infect Dis, 191, 1771–7.

